# Punchline: Identifying and comparing significant Pfam protein domain differences across draft whole genome sequences

**DOI:** 10.1101/686543

**Authors:** Lisa C Crossman

## Abstract

**Motivation:** Short-read draft paired-end Illumina assemblies can be fragmented, contain many contigs and be impacted on by repeat regions, caused by mobile element activity within the genome or inherently repetitive gene structure. Annotating such assemblies for function and analysing gene content can be challenging if predicted genes are fragmented across contigs. Such a case can often occur within specific families of genes such as longer genes with repeating domains, genes specifying several transmembrane domains and of unusual nucleotide content. These genes can often be virulence determinants, therefore losing these specific types of data can seriously impact downstream studies.

**Results:** Rather than studying the predicted gene content of draft genomes, we examined predicted protein content using the Pfam domain complements of predicted proteins. We produced a workflow, Punchline, to study the genetic content of draft contig assemblies by looking at the complement of short domains that are unlikely to be affected. We investigated a dataset of *Bacteroides ovatus* in terms of a grouping involving the vertebrate host from which the organism was isolated and identified potential host restricted functions and host restricted phylogenetic clustering.

**Availability:** https://github.com/LCrossman

## 1 Introduction

Whole genome assemblies from short-read sequencing data are subject to factors that impact on their quality including repeats and regions of poor sequence coverage. Very high and low % GC genomes may present with additional technical issues. These impacts impinge on short-read draft whole genome assemblies and can result in fragmentation across these regions.

Gene prediction programs can struggle in annotating draft assemblies with several short contigs, and larger genes, particularly those with repeats that can be broken across contigs. Down-stream studies involving looking at the presence/absence of specific genes under different conditions or pan-core genome analyses can be particularly affected by high levels of assembly fragmentation. Unfortunately, membrane-bound genes with repeating domains can be of high relevance as species-specific or isolate-specific genes of interest. In addition, genes encoding surface proteins/ antigens can fall into this category. These genes can also be of particular interest in studies of pathogenicity and should be considered in downstream investigations.

To lessen the impact of fragmented assemblies on the preence/absence of specific genes, this study looks at the statistical importance of protein family domains identified from the Protein families database, Pfam (Finn *et al.*, 2014; El-Gebali *et al.*, 2018). The underlying premise of this study is that whilst whole genes may be fragmented and broken across contigs, they are much less likely to have fragmented Pfam domains. Importantly for pangenome studies, genes may share large regions of similarity but have specific smaller regions of difference that can be detected by the examination of predicted Pfam domain complements but may be missed by studies of entire gene complement. Pfam domains are detected using Hidden Markov Models (HMM), a machine learning technique where the models are built from training examples which are known good members (Eddy, 2011; Mistry *et al.*, 2013). The speed of detection of Pfam domains using HMMER is a distinct advantage to BLAST-based methods of looking at gene orthologues for whole core and accessory genes. This includes an improved detection method due to the immediate provision of functional information which improves an overall understanding of function.

In this study, we provide a highly scalable workflow to detect numbers and types of Pfam domains within predicted protein datasets for fragmented draft assemblies. We identify significant Pfam domains between specific metadata groups using statistical significance testing. In addition, a distance matrix is built from the counted numbers of the Pfam domain complement of each genome. This allows a distance metric to be produced using normalised domain counts that can be used to draw phylogenetic trees of relatedness. Correspondingly, we provide a means of correlating top Pfam domains, showing which are found in association together and provide means to visualise the data.

## 2 Methods

The workflow is tested compatible with Python 2.7, 3.5, 3.7 and R 3.3.1, 3.4.1, 3.5.0 (R Development Core Team, 2015, 2017). Required dependencies are HMMER3 (Eddy, 2011) and the Pfam A database (Finn *et al.*, 2014; El-Gebali *et al.*, 2018). The R libraries required are ape (Paradis *et al.*, 2004), optparse (Davis, 2019), gplots (Warnes *et al.*, 2016), ggplot2 (Wickham, 2016) and corrplot (Wei and Simko, 2016) and for phylogenetic clustering, Lsa (Wild, 2007), each available from Bioconductor (Huber *et al.*, 2015). Phylip is a requirement for the phylogenetic clustering (Felsenstein, 2005). In the workflow, draft genome assemblies as genbank or embl files are required as input or a proteome as protein fasta file such as can be downloaded directly from Uniprot proteomes (Bateman, 2019). On each, a protein fasta dataset is built then a Pfam search is run with HMMER3 in parallel with specific parameters to produce a domain table. From the results of running a Pfam search on each assembly, the table is combined, and raw counts produced. The raw counts can be compared as they stand, or compared with normalization using either relative or log-relative transformation to account for differences in Pfam domain raw counts. For example, it may be required to compare shorter and longer genomes. From the domain counts, a distance metric is calculated, and a phylogenetic tree can be drawn. In addition, the numbers of domains can be used in a correlation matrix to produce a heatmap showing which domains are found in association with which other domains. If a grouping file is provided, statistics can be produced for significant domain differences between the groups using the Kruskal-Wallis test, also termed the one-way ANOVA on ranks or one-way analysis of variance, (MacDonald, 2009) with the Benjamini-Hochberg correction for multiple testing (Ferreira and Zwinderman, 2006).

## 3 Results

**3.1** Workflow

The steps employed in the workflow are as follows:

1. Input genome assemblies as protein fasta files and if required, a grouping file
2. Run HMMER3 against Pfam A in parallel
3. Collect results and combining separate Pfam tables into a matrix
4. Kruskal-Wallis statistical significance testing with specific groups
5. Correlation heatmap plot with Pfam domains correlated with other Pfam domains discovered present.
6. Phylogenetic clustering from distance metric calculated using the Pfam domain counts

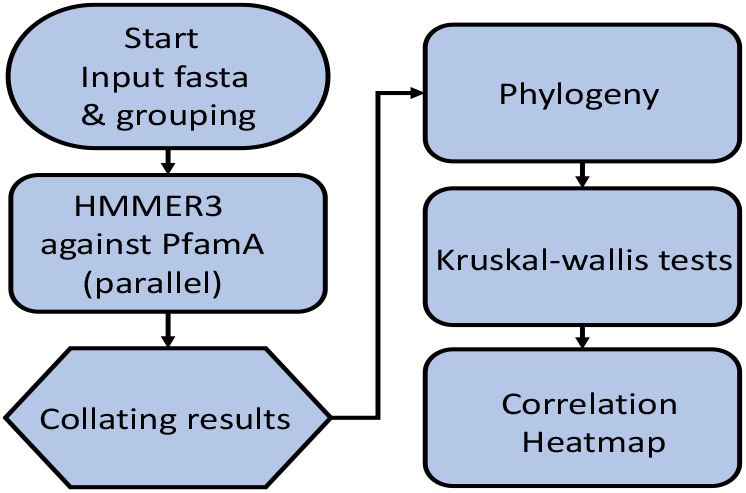

Once the workflow stages have been carried out, the outputs of the process are the following:

- Statistical tables of numbers of Pfam domains found significantly higher in one group category over the other.
- Heatmap data of the top Pfam domains by sample.
- Correlation heatmap matrix of Pfam domain associations.
- Phylogenetic tree with branches coloured by group category.

HMMER3 has the advantage of high scalability and a low running time with the requirement for minimal compute resources (Eddy, 2011).

### 3.1 Specific Application example

A test dataset comprising draft assemblies of whole genome sequences of *Bacteroides ovatus* from the sequence databases were investigated with the workflow (Supplementary table 1). These were isolated from alternative vertebrate gut host niches, in particular, bovine, hominine (human), porcine and caprine (goat).

*B. ovatus* has been described the Swiss-army knife of the digestive tract, due to its unusual ability to carry out xyloglucan (XyG) degradation. XyG is a hemicellulose found in plant cell walls. Despite XyG being an important part of the human diet, we do not possess enzymes to degrade it. The degradation machinery is found specific to particular species of Bacteroidetes (Larsbrink *et al.*, 2014). The prevalence of XyGs in the human diet suggests that the mechanism by which bacteria degrade these complex polysaccharides is highly important to human energy acquisition. The rarity of XyG metabolism highlights the significance of *B. ovatus* and other proficient XyG degraders as key members of the human gut microbial consortium (Larsbrink *et al.*, 2014; Flint *et al.*, 2012).

Full significance testing results using Kruskal-wallis tests are presented in Supplementary table 2. In addition to the previous analyses, phylogenetic clustering was performed by first building a cosine similarity matrix using latent semantic analysis (lsa) from Pfam domain counts as fingerprints of the strains of both *B. ovatus* and *B. thetaiotaomicron*. A distance matrix was created from the similarity matrix then a neighbor joining tree was performed using Phylip neighbor (Figure 4). Finally, the tree was plotted using the R ape package. These data indicate the potential for some host restricted functions whilst phylogenetic clustering suggests some host restricted clades. Some of these functions are shown in Supplementary table 3 with boxplot data in Supplementary Figure 1. Host restricted clades have been seen previously amongst *Lactobacillus reuteri* (Frese *et al.*, 2011; Wegmann *et al.*, 2015). Reasons for seeing isolates with alternative specificities in different host niches could be down to niche specialization, partial niche overlap (Wexler and Goodman, 2017) or alternative dietary specificity.

**Figure 2.**
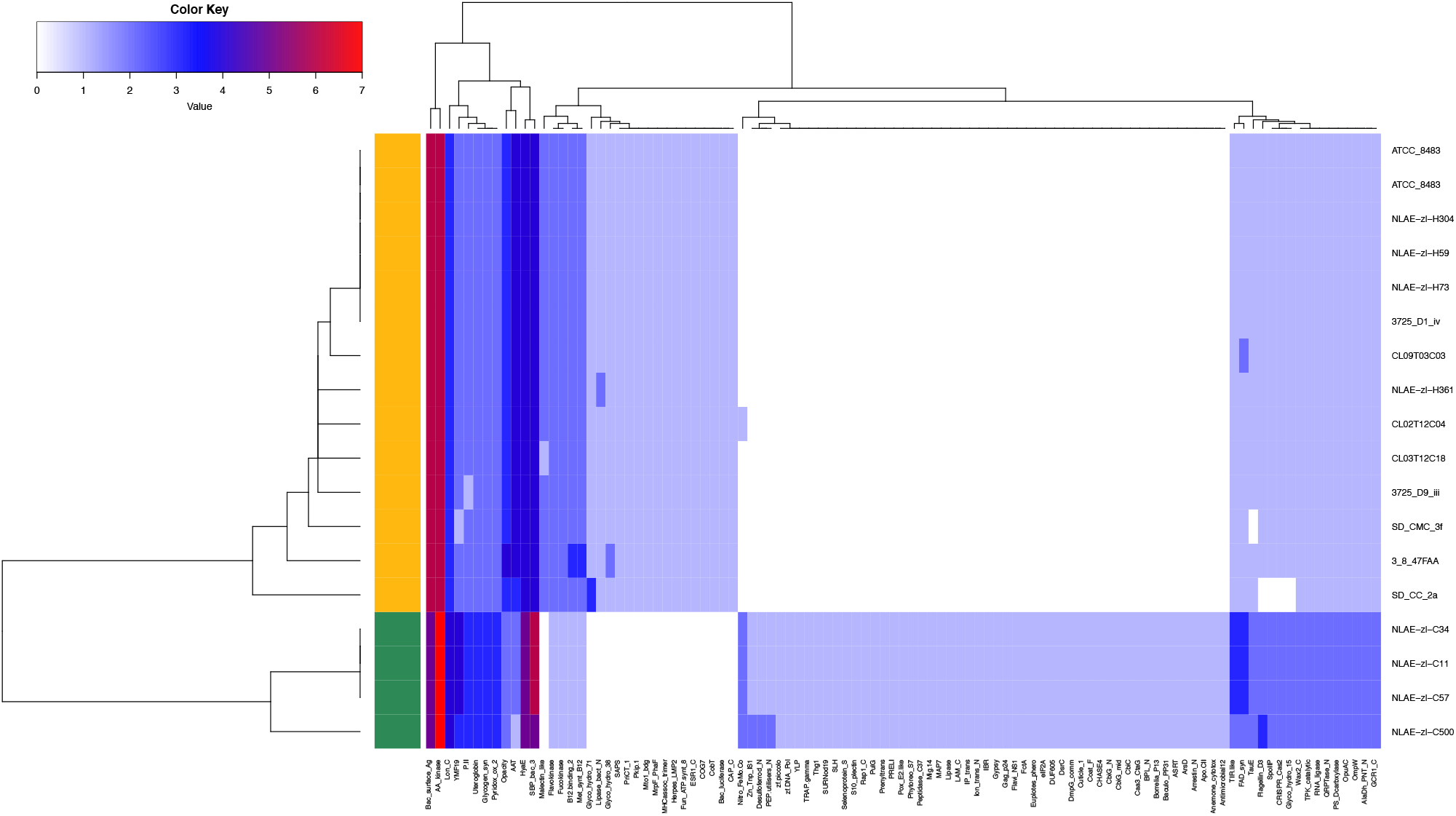
Heatmap of *B. ovatus* Pfam domain counts showing significant differences across categories (alternative vertebrate hosts) In the heatmap figure above, Pfam domain counts for predicted proteins in *B. ovatus* isolated from either human (gold) or bovine (green) hosts. Counts are denoted using colour palette migrating from low (white) to mid (blue) and high (red) counts. Clustering of isolates is shown as a dendrogram in the row dimension whilst clustering of Pfam domain counts are shown as a dendrogram in the column dimension. We can clearly see sets of Pfam domains that are shared among particular isolates.

**Figure 3.**
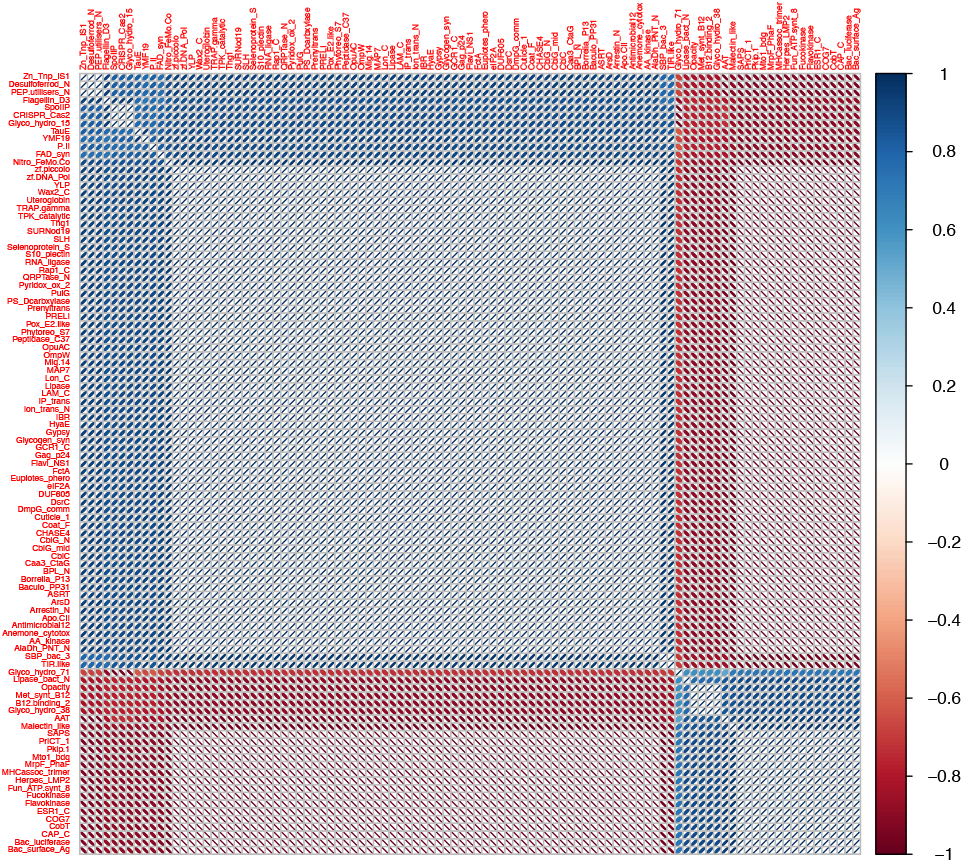
Correlation plot of Pfam domain counts. In the correlation plot above, blue ellipses show positive correlation whilst red ellipses show negative correlation scores. The more the ellipse tends towards a circle, the weaker the correlation. We can clearly see at least two groups of Pfam domain that can be found together and correlate negatively with the alternative group. The domains may be found within the same protein or the same metabolic pathway or may be functionally associated in another manner.

**Figure 4.**
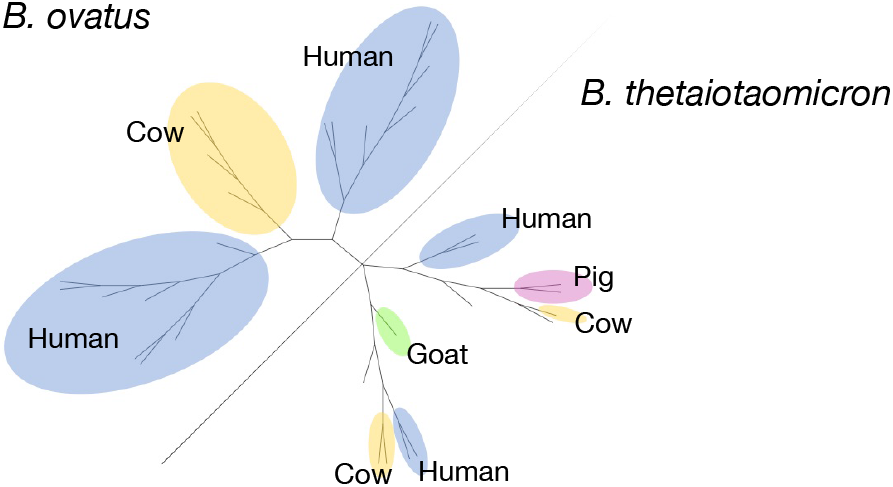
Clustering of Pfam domain counts of exemplar genomes. Phylogenetic clustering using latent semantic analysis of Pfam domain counts to create first a cosine similarity matrix, from that a distance matrix and a neighbor joining tree. The tree was visualized unrooted using R ape package and modified to include metadata. Host restricted clades are suggested. There was also some evidence of geographical clustering, which was more pronounced among the *B.thetaiotaomicron* dataset.

## 4 Discussion

The applications of the process lie firstly in examining functional information from fragmented genome assemblies. The outputs provide a means by which to study different functional domains across different groups of strains such that fragmented genome assemblies do not appreciably impact on the results. A phylogenetic fingerprint provides a tree grouped by functional domain information and a correlation heatmap can be used to study high level associations between domains. The information provides the analyst with functional data beyond a simple gene presence or absence. Correlations of the Pfam domains can indicate functions across the genomes.

Whilst Pfam data is extremely informative in terms of producing functional data, it is of course important to note that some important genes or virulence determinants do not possess characterised Pfam domains.

On the other hand, particular classes of genes can be lost from core and accessory studies of genomic gene complements due to their length and unusual composition. Directly studying the Pfam domain counts can provide us with a means to counteract losing the information from genes fragmented across assemblies whilst also providing a set of useful functional data. The speed of Pfam searches *via* HMMER3 and of this workflow means that it is highly scalable to many genomes and results give instant functional information without requiring lengthy studies of specific genes.

## Acknowledgements

We thank Dr. Gemma Langridge, Quadram Institute for helpful comments.

## Conflict of Interest

none declared.

---

**Supplementary table 1:**

Accession numbers of genomes used in the example study

**Supplementary table 2:**

Significance tests of Pfam domain count differences across the predicted proteins from the exemplar draft whole genome sequences.

**Supplementary table 3:**

Suggested functions of significant Pfam domains of interest

**Supplementary Figure 1:**

Boxplot graph of significant Pfam domains of *B. ovatus* in bovine compared to human hosts

