## Supplementary table 2 for "Punchline: Identifying and comparing significant Pfam protein domain differences across draft whole genome sequences"

| id | p.value | E.value | FWER | q.value |
|---|---|---|---|---|
| Zn_Tnp_IS1 | 4.15E-05 | 0.198529188 | 60.50632911 | 0.002513028 |
| TIR-like | 4.15E-05 | 0.198529188 | 61.28205128 | 0.002545246 |
| PulG | 4.15E-05 | 0.198529188 | 62.07792208 | 0.002578301 |
| zf-piccolo | 3.74E-05 | 0.178675532 | 69.27536232 | 0.0025895 |
| PS_Dcarbxylase | 4.15E-05 | 0.198529188 | 62.89473684 | 0.002612226 |
| zf-DNA_Pol | 3.74E-05 | 3.74E-05 | 70.29411765 | 0.002627581 |
| PEP-utilisers_N | 4.15E-05 | 0.198529188 | 63.73333333 | 0.002647056 |
| YLP | 3.74E-05 | 0.178675532 | 71.34328358 | 0.002666799 |
| Lon_C | 4.15E-05 | 4.15E-05 | 64.59459459 | 0.002682827 |
| Wax2_C | 3.74E-05 | 0.178675532 | 72.42424242 | 0.002707205 |
| Ion_trans_N | 4.15E-05 | 4.15E-05 | 65.47945205 | 0.002719578 |
| TRAP-gamma | 3.74E-05 | 3.74E-05 | 73.53846154 | 0.002748854 |
| HlyU | 4.15E-05 | 0.198529188 | 66.38888889 | 0.00275735 |
| TPK_catalytic | 3.74E-05 | 3.74E-05 | 74.6875 | 0.002791805 |
| Glycogen_syn | 4.15E-05 | 4.15E-05 | 67.32394366 | 0.002796186 |
| Thg1 | 3.74E-05 | 0.178675532 | 75.87301587 | 0.00283612 |
| Desulfoferrod_N | 4.15E-05 | 0.198529188 | 68.28571429 | 0.002836131 |
| SURNod19 | 3.74E-05 | 3.74E-05 | 77.09677419 | 0.002881863 |
| SLH | 3.74E-05 | 0.178675532 | 78.36065574 | 0.002929107 |
| Selenoprotein_S | 3.74E-05 | 0.178675532 | 79.66666667 | 0.002977926 |
| SAPS | 3.74E-05 | 3.74E-05 | 81.01694915 | 0.003028399 |
| S10_plectin | 3.74E-05 | 3.74E-05 | 82.4137931 | 0.003080613 |
| RNA_ligase | 3.74E-05 | 0.178675532 | 83.85964912 | 0.003134658 |
| Rap1_C | 3.74E-05 | 0.178675532 | 85.35714286 | 0.003190634 |
| QRPTase_N | 3.74E-05 | 0.178675532 | 86.90909091 | 0.003248646 |
| Pyridox_ox_2 | 3.74E-05 | 3.74E-05 | 88.51851852 | 0.003308806 |
| Prenyltrans | 3.74E-05 | 0.178675532 | 90.18867925 | 0.003371236 |
| PRELI | 3.74E-05 | 3.74E-05 | 91.92307692 | 0.003436068 |
| Pox_E2-like | 3.74E-05 | 3.74E-05 | 93.7254902 | 0.003503442 |
| Pkip-1 | 3.74E-05 | 3.74E-05 | 95.6 | 0.003573511 |
| Phytoreo_S7 | 3.74E-05 | 0.178675532 | 97.55102041 | 0.003646439 |
| Peptidase_C37 | 3.74E-05 | 3.74E-05 | 99.58333333 | 0.003722407 |
| OpuAC | 3.74E-05 | 3.74E-05 | 101.7021277 | 0.003801607 |
| OmpW | 3.74E-05 | 0.178675532 | 103.9130435 | 0.003884251 |
| Mto1_bdg | 3.74E-05 | 3.74E-05 | 106.2222222 | 0.003970567 |
| Mig-14 | 3.74E-05 | 3.74E-05 | 108.6363636 | 0.004060808 |
| MHCassoc_trimer | 3.74E-05 | 3.74E-05 | 111.1627907 | 0.004155245 |
| MAP7 | 3.74E-05 | 3.74E-05 | 113.8095238 | 0.004254179 |
| Lipase | 3.74E-05 | 0.178675532 | 116.5853659 | 0.00435794 |
| LAM_C | 3.74E-05 | 0.178675532 | 119.5 | 0.004466888 |
| IP_trans | 3.74E-05 | 3.74E-05 | 122.5641026 | 0.004581424 |
| IBR | 3.74E-05 | 3.74E-05 | 125.7894737 | 0.004701988 |
| HyaE | 3.74E-05 | 3.74E-05 | 129.1891892 | 0.004829068 |
| Herpes_LMP2 | 3.74E-05 | 0.178675532 | 132.7777778 | 0.004963209 |
| Gypsy | 3.74E-05 | 0.178675532 | 136.5714286 | 0.005105015 |

| | | | | |
|---|---|---|---|---|
| Gag_p24 | 3.74E-05 | 3.74E-05 | 140.5882353 | 0.005255163 |
| Fun_ATP-synt_8 | 3.74E-05 | 3.74E-05 | 144.8484848 | 0.00541441 |
| Fucokinase | 3.74E-05 | 3.74E-05 | 149.375 | 0.00558361 |
| Flavokinase | 3.74E-05 | 3.74E-05 | 154.1935484 | 0.005763727 |
| Flavi_NS1 | 3.74E-05 | 3.74E-05 | 159.3333333 | 0.005955851 |
| FctA | 3.74E-05 | 3.74E-05 | 164.8275862 | 0.006161225 |
| Euplotes_phero | 3.74E-05 | 3.74E-05 | 170.7142857 | 0.006381269 |
| eIF2A | 3.74E-05 | 3.74E-05 | 177.037037 | 0.006617612 |
| DUF605 | 3.74E-05 | 0.178675532 | 183.8461538 | 0.006872136 |
| DsrC | 3.74E-05 | 0.178675532 | 191.2 | 0.007147021 |
| WD40 | 0.000147802 | 0.706495043 | 49.27835052 | 0.007283454 |
| Uteroglobin | 0.000147802 | 0.706495043 | 49.79166667 | 0.007359323 |
| TauE | 0.000147802 | 0.706495043 | 50.31578947 | 0.00743679 |
| DmpG_comm | 3.74E-05 | 3.74E-05 | 199.1666667 | 0.007444814 |
| SpoIIP | 0.000147802 | 0.706495043 | 50.85106383 | 0.007515905 |
| Rabaptin | 0.000147802 | 0.000147802 | 51.39784946 | 0.007596721 |
| YMF19 | 0.000159531 | 0.000159531 | 47.8 | 0.007625584 |
| PriCT_1 | 0.000147802 | 0.000147802 | 51.95652174 | 0.007679294 |
| Flagellin_D3 | 0.000159531 | 0.000159531 | 48.28282828 | 0.00770261 |
| P-II | 0.000147802 | 0.000147802 | 52.52747253 | 0.007763682 |
| Cuticle_1 | 3.74E-05 | 3.74E-05 | 207.826087 | 0.007768501 |
| AAT | 0.000159531 | 0.762558369 | 48.7755102 | 0.007781208 |
| Nitro_FeMo-Co | 0.000147802 | 0.000147802 | 53.11111111 | 0.007849945 |
| MrpF_PhaF | 0.000147802 | 0.000147802 | 53.70786517 | 0.007938147 |
| Met_synt_B12 | 0.000147802 | 0.706495043 | 54.31818182 | 0.008028353 |
| Lipase_bact_N | 0.000147802 | 0.706495043 | 54.94252874 | 0.008120633 |
| COG7 | 3.74E-05 | 3.74E-05 | 217.2727273 | 0.008121615 |
| Glyco_hydro_71 | 0.000147802 | 0.000147802 | 55.58139535 | 0.008215059 |
| Glyco_hydro_38 | 0.000147802 | 0.000147802 | 56.23529412 | 0.008311706 |
| Glyco_hydro_15 | 0.000147802 | 0.706495043 | 56.9047619 | 0.008410655 |
| CobT | 3.74E-05 | 0.178675532 | 227.6190476 | 0.008508359 |
| GCR1_C | 0.000147802 | 0.706495043 | 57.59036145 | 0.008511988 |
| CRISPR_Cas2 | 0.000147802 | 0.706495043 | 58.29268293 | 0.008615793 |
| C1_2 | 0.000147802 | 0.000147802 | 59.01234568 | 0.008722161 |
| Bac_surface_Ag | 0.000147802 | 0.000147802 | 59.75 | 0.008831188 |
| Coat_F | 3.74E-05 | 0.178675532 | 239 | 0.008933777 |
| CHASE4 | 3.74E-05 | 3.74E-05 | 251.5789474 | 0.009403975 |
| FAD_syn | 0.000206109 | 0.000206109 | 46.8627451 | 0.009658839 |
| FAD_binding_4 | 0.000206109 | 0.000206109 | 47.32673267 | 0.009754471 |
| CbiG_N | 3.74E-05 | 0.178675532 | 265.5555556 | 0.009926418 |
| CbiG_mid | 3.74E-05 | 3.74E-05 | 281.1764706 | 0.010510325 |
| CbiC | 3.74E-05 | 0.178675532 | 298.75 | 0.011167221 |
| CAP_C | 3.74E-05 | 3.74E-05 | 318.6666667 | 0.011911702 |
| zf-met2 | 0.000561339 | 2.68320253 | 21.62895928 | 0.012141188 |
| YjgP_YjgQ | 0.000561339 | 2.68320253 | 21.72727273 | 0.012196375 |
| VirJ | 0.000561339 | 0.000561339 | 21.82648402 | 0.012252066 |

| | | | | |
|---|---|---|---|---|
| Urm1 | 0.000561339 | 0.000561339 | 21.9266055 | 0.012308268 |
| UAA | 0.000561339 | 0.000561339 | 22.02764977 | 0.012364989 |
| TUDOR | 0.000561339 | 2.68320253 | 22.12962963 | 0.012422234 |
| Toxin_28 | 0.000561339 | 0.000561339 | 22.23255814 | 0.012480012 |
| Tom37_C | 0.000561339 | 0.000561339 | 22.3364486 | 0.01253833 |
| FTH | 0.000591399 | 2.826886816 | 21.24444444 | 0.012563941 |
| ThiW | 0.000561339 | 0.000561339 | 22.44131455 | 0.012597195 |
| TRAM1 | 0.000588344 | 0.000588344 | 21.43497758 | 0.012611136 |
| CbiJ | 0.000591399 | 0.000591399 | 21.33928571 | 0.01262003 |
| TFCD_C | 0.000561339 | 2.68320253 | 22.54716981 | 0.012656616 |
| MukB | 0.000588344 | 0.000588344 | 21.53153153 | 0.012667943 |
| Syndecan | 0.000561339 | 0.000561339 | 22.65402844 | 0.0127166 |
| Caa3_CtaG | 3.74E-05 | 0.178675532 | 341.4285714 | 0.012762538 |
| SpoU_methylase | 0.000561339 | 0.000561339 | 22.76190476 | 0.012777155 |
| SLIDE | 0.000561339 | 0.000561339 | 22.8708134 | 0.01283829 |
| WD-3 | 0.000379895 | 0.000379895 | 33.90070922 | 0.012878723 |
| She2p | 0.000561339 | 0.000561339 | 22.98076923 | 0.012900012 |
| SH3_7 | 0.000561339 | 2.68320253 | 23.09178744 | 0.012962331 |
| VanW | 0.000379895 | 1.815899935 | 34.14285714 | 0.012970714 |
| SCHIP-1 | 0.000561339 | 2.68320253 | 23.2038835 | 0.013025255 |
| TryThrA_C | 0.000379895 | 1.815899935 | 34.38848921 | 0.013064028 |
| SbcD_C | 0.000561339 | 2.68320253 | 23.31707317 | 0.013088793 |
| Orbi_NS3 | 0.000389584 | 0.000389584 | 33.66197183 | 0.013114179 |
| Tenui_NS3 | 0.000639565 | 0.000639565 | 20.51502146 | 0.013120688 |
| SAYSvFN | 0.000561339 | 2.68320253 | 23.43137255 | 0.013152954 |
| Transposase_30 | 0.000379895 | 1.815899935 | 34.63768116 | 0.013158695 |
| MIP | 0.000639565 | 0.000639565 | 20.60344828 | 0.013177243 |
| RST | 0.000561339 | 0.000561339 | 23.54679803 | 0.013217746 |
| Malectin | 0.000639565 | 3.057120293 | 20.69264069 | 0.013234287 |
| Toxin_R_bind_N | 0.000379895 | 0.000379895 | 34.89051095 | 0.013254744 |
| Rrn6 | 0.000561339 | 0.000561339 | 23.66336634 | 0.013283181 |
| Trehalose_PPase | 0.000656068 | 3.136002925 | 20.25423729 | 0.013288148 |
| Glyco_hydro_57 | 0.000639565 | 3.057120293 | 20.7826087 | 0.013291827 |
| URO-D | 0.000661633 | 3.162606045 | 20.16877637 | 0.013344329 |
| SH3_2 | 0.000656068 | 0.000656068 | 20.34042553 | 0.013344693 |
| Rod_C | 0.000561339 | 2.68320253 | 23.78109453 | 0.013349266 |
| GcrA | 0.000639565 | 0.000639565 | 20.87336245 | 0.01334987 |
| Toxin_R_bind_C | 0.000379895 | 1.815899935 | 35.14705882 | 0.013352205 |
| Acyltransferase | 0.000656068 | 0.000656068 | 20.42735043 | 0.013401722 |
| ELFV_dehydrog | 0.000639565 | 3.057120293 | 20.96491228 | 0.013408422 |
| RNA_bind | 0.000561339 | 2.68320253 | 23.9 | 0.013416013 |
| REV | 0.000379895 | 0.000379895 | 35.40740741 | 0.013451111 |
| eIF3_p135 | 0.000639565 | 3.057120293 | 21.05726872 | 0.01346749 |
| RL11D | 0.000561339 | 2.68320253 | 24.0201005 | 0.01348343 |
| GDI | 0.000672862 | 3.216278108 | 20.08403361 | 0.013513774 |
| Alpha-mann_mid | 0.000639565 | 0.000639565 | 21.15044248 | 0.013527081 |

| | | | | |
|---|---|---|---|---|
| Peptidase_S28 | 0.000379895 | 1.815899935 | 35.67164179 | 0.013551492 |
| Rif1_N | 0.000561339 | 0.000561339 | 24.14141414 | 0.013551528 |
| Ribosomal_L31 | 0.000561339 | 0.000561339 | 24.26395939 | 0.013620317 |
| Orthopox_35kD | 0.000379895 | 1.815899935 | 35.93984962 | 0.013653383 |
| TP_methylase | 0.000689949 | 3.297956865 | 19.8340249 | 0.013684468 |
| PseudoU_synth_1 | 0.000561339 | 2.68320253 | 24.3877551 | 0.013689809 |
| Nudix_N | 0.000689949 | 0.000689949 | 19.91666667 | 0.013741487 |
| BPL_N | 3.74E-05 | 3.74E-05 | 367.6923077 | 0.013744272 |
| Nit_Regul_Hom | 0.000379895 | 1.815899935 | 36.21212121 | 0.013756818 |
| PRRSV_Env | 0.000561339 | 2.68320253 | 24.51282051 | 0.013760013 |
| CbiG_C | 0.000689949 | 3.297956865 | 20 | 0.013798983 |
| Pox_F16 | 0.000561339 | 2.68320253 | 24.63917526 | 0.013830941 |
| NiFeSe_Hases | 0.000379895 | 0.000379895 | 36.48854962 | 0.013861832 |
| PM0188 | 0.000561339 | 0.000561339 | 24.76683938 | 0.013902604 |
| Menin | 0.000379895 | 1.815899935 | 36.76923077 | 0.013968461 |
| PLDc | 0.000561339 | 0.000561339 | 24.89583333 | 0.013975013 |
| Pico_P2B | 0.000561339 | 0.000561339 | 25.02617801 | 0.014048181 |
| Lipoprotein_10 | 0.000379895 | 1.815899935 | 37.05426357 | 0.014076744 |
| Phage_Cox | 0.000561339 | 0.000561339 | 25.15789474 | 0.014122119 |
| Glypican | 0.000379895 | 0.000379895 | 37.34375 | 0.014186718 |
| Pep3_Vps18 | 0.000561339 | 0.000561339 | 25.29100529 | 0.014196839 |
| p450 | 0.000561339 | 2.68320253 | 25.42553191 | 0.014272354 |
| Gamma-COP | 0.000379895 | 0.000379895 | 37.63779528 | 0.014298425 |
| OrfB_IS605 | 0.000561339 | 2.68320253 | 25.56149733 | 0.014348677 |
| Fusion_gly | 0.000379895 | 1.815899935 | 37.93650794 | 0.014411904 |
| Orbi_VP4 | 0.000561339 | 2.68320253 | 25.69892473 | 0.01442582 |
| NapD | 0.000561339 | 2.68320253 | 25.83783784 | 0.014503797 |
| Fimbrial | 0.000379895 | 1.815899935 | 38.24 | 0.014527199 |
| PC_rep | 0.000740823 | 3.541132556 | 19.67078189 | 0.014572562 |
| MRP-S35 | 0.000561339 | 2.68320253 | 25.97826087 | 0.014582622 |
| Arf | 0.000740823 | 3.541132556 | 19.75206612 | 0.014632779 |
| DUF3882 | 0.000379895 | 1.815899935 | 38.5483871 | 0.014644354 |
| MnhB | 0.000561339 | 0.000561339 | 26.12021858 | 0.014662309 |
| Malate_synthase | 0.000561339 | 0.000561339 | 26.26373626 | 0.014742871 |
| DpnII | 0.000379895 | 1.815899935 | 38.86178862 | 0.014763414 |
| Lyase_8_N | 0.000561339 | 0.000561339 | 26.40883978 | 0.014824323 |
| Cytochrom_B561 | 0.000379895 | 0.000379895 | 39.18032787 | 0.014884426 |
| BphX | 3.74E-05 | 3.74E-05 | 398.3333333 | 0.014889628 |
| Lipoprotein_18 | 0.000561339 | 0.000561339 | 26.55555556 | 0.014906681 |
| Opacity | 0.000359634 | 0.000359634 | 41.56521739 | 0.01494825 |
| KaiA | 0.000561339 | 0.000561339 | 26.70391061 | 0.014989958 |
| COX_ARM | 0.000379895 | 1.815899935 | 39.50413223 | 0.015007437 |
| UXS1_N | 0.000356009 | 1.7017238 | 42.30088496 | 0.015059503 |
| Hum_adeno_E3A | 0.000561339 | 0.000561339 | 26.85393258 | 0.015074172 |
| Cytochrom_C_2 | 0.000359634 | 0.000359634 | 41.92982456 | 0.015079375 |
| C2 | 0.000379895 | 0.000379895 | 39.83333333 | 0.015132499 |

| | | | | |
|---|---|---|---|---|
| HTH_9 | 0.000561339 | 2.68320253 | 27.00564972 | 0.015159336 |
| RDD | 0.000356009 | 1.7017238 | 42.67857143 | 0.015193963 |
| HPPK | 0.000561339 | 2.68320253 | 27.15909091 | 0.015245469 |
| Adeno_IVa2 | 0.000379895 | 0.000379895 | 40.16806723 | 0.015259663 |
| Aminoglyc_resit | 0.000328851 | 1.571907551 | 46.40776699 | 0.015261238 |
| Pyridox_oxidase | 0.000356009 | 1.7017238 | 43.06306306 | 0.015330845 |
| HobA | 0.000561339 | 0.000561339 | 27.31428571 | 0.015332586 |
| AA_permease | 0.000379895 | 1.815899935 | 40.50847458 | 0.015388982 |
| Hira | 0.000561339 | 2.68320253 | 27.47126437 | 0.015420704 |
| PuR_N | 0.000356009 | 1.7017238 | 43.45454545 | 0.015470216 |
| Hexokinase_1 | 0.000561339 | 0.000561339 | 27.6300578 | 0.015509841 |
| 5_3_exonuc_N | 0.000379895 | 0.000379895 | 40.85470085 | 0.015520512 |
| GDE_C | 0.000378152 | 1.807568363 | 41.20689655 | 0.015582486 |
| HEAT_PBS | 0.000561339 | 2.68320253 | 27.79069767 | 0.015600015 |
| FtsL | 0.000356009 | 0.000356009 | 43.85321101 | 0.015612145 |
| HCV_NS4b | 0.000561339 | 0.000561339 | 27.95321637 | 0.015691243 |
| ESR1_C | 0.000356009 | 1.7017238 | 44.25925926 | 0.015756702 |
| GcpE | 0.000561339 | 0.000561339 | 28.11764706 | 0.015783544 |
| Flt3_lig | 0.000561339 | 2.68320253 | 28.28402367 | 0.015876938 |
| Col_cuticle_N | 0.000356009 | 0.000356009 | 44.6728972 | 0.015903961 |
| Flo11 | 0.000561339 | 2.68320253 | 28.45238095 | 0.015971444 |
| Colicin_im | 0.000356009 | 1.7017238 | 45.09433962 | 0.016053998 |
| FlaE | 0.000561339 | 0.000561339 | 28.62275449 | 0.016067081 |
| FAD-oxidase_C | 0.000561339 | 2.68320253 | 28.79518072 | 0.016163871 |
| CbiX | 0.000356009 | 1.7017238 | 45.52380952 | 0.016206893 |
| Borrelia_P13 | 3.74E-05 | 3.74E-05 | 434.5454545 | 0.01624323 |
| F-actin_cap_A | 0.000561339 | 2.68320253 | 28.96969697 | 0.016261834 |
| EXS | 0.000561339 | 2.68320253 | 29.14634146 | 0.016360991 |
| B12-binding_2 | 0.000356009 | 1.7017238 | 45.96153846 | 0.016362729 |
| PmbA_TldD | 0.000843559 | 4.032210211 | 19.43089431 | 0.016391098 |
| NdhM | 0.000843559 | 4.032210211 | 19.51020408 | 0.016458001 |
| Endonuc_Holl | 0.000561339 | 0.000561339 | 29.32515337 | 0.016461365 |
| EcsC | 0.000843559 | 0.000843559 | 19.59016393 | 0.016525452 |
| ELL | 0.000561339 | 2.68320253 | 29.50617284 | 0.016562979 |
| EF1_GNE | 0.000561339 | 0.000561339 | 29.68944099 | 0.016665854 |
| EB1 | 0.000561339 | 0.000561339 | 29.875 | 0.016770016 |
| E3_UbLigase_EDD | 0.000561339 | 0.000561339 | 30.06289308 | 0.016875488 |
| DRY_EERY | 0.000561339 | 2.68320253 | 30.25316456 | 0.016982294 |
| DNA_pol_alpha_N | 0.000561339 | 0.000561339 | 30.44585987 | 0.017090462 |
| DBR1 | 0.000561339 | 0.000561339 | 30.64102564 | 0.017200016 |
| DASH_Ask1 | 0.000561339 | 0.000561339 | 30.83870968 | 0.017310984 |
| CofC | 0.000561339 | 2.68320253 | 31.03896104 | 0.017423393 |
| CENP-M | 0.000561339 | 0.000561339 | 31.24183007 | 0.017537271 |
| Cbl_N3 | 0.000561339 | 0.000561339 | 31.44736842 | 0.017652648 |
| Cas_Cas02710 | 0.000561339 | 0.000561339 | 31.65562914 | 0.017769553 |
| Transposase_22 | 0.000946925 | 4.526300726 | 18.81889764 | 0.017820082 |

| | | | | |
|---|---|---|---|---|
| Bac_luciferase | 3.74E-05 | 3.74E-05 | 478 | 0.017867553 |
| Calmodulin_bind | 0.000561339 | 2.68320253 | 31.86666667 | 0.017888017 |
| POTRA_2 | 0.000946925 | 4.526300726 | 18.89328063 | 0.017890517 |
| Mif2 | 0.000946925 | 4.526300726 | 18.96825397 | 0.017961511 |
| Calci_bind_CcbP | 0.000561339 | 2.68320253 | 32.08053691 | 0.018008071 |
| DNA_ligase_ZBD | 0.000946925 | 4.526300726 | 19.0438247 | 0.018033071 |
| CoA_binding | 0.000946925 | 0.000946925 | 19.12 | 0.018105203 |
| Bunya_RdRp | 0.000561339 | 2.68320253 | 32.2972973 | 0.018129747 |
| CBAH | 0.000946925 | 4.526300726 | 19.19678715 | 0.018177915 |
| ketoacyl-synt | 0.0009446 | 0.0009446 | 19.27419355 | 0.01820641 |
| Bacteriocin_IIi | 0.000561339 | 0.000561339 | 32.5170068 | 0.018253078 |
| Acatn | 0.0009446 | 4.515189557 | 19.35222672 | 0.01828012 |
| Cas_APE2256 | 0.000548423 | 2.621460427 | 33.42657343 | 0.018331891 |
| SBP_bac_3 | 0.000990324 | 0.000990324 | 18.52713178 | 0.018347865 |
| Arv1 | 0.000561339 | 2.68320253 | 32.73972603 | 0.0183781 |
| Rota_NSP4 | 0.000990324 | 0.000990324 | 18.59922179 | 0.018419258 |
| Inhibitor_I48 | 0.000990324 | 4.733749251 | 18.671875 | 0.018491208 |
| APP_E2 | 0.000561339 | 2.68320253 | 32.96551724 | 0.018504845 |
| NAD_binding_3 | 0.001005081 | 0.001005081 | 18.45559846 | 0.018549379 |
| GFO_IDH_MocA_C | 0.000990324 | 4.733749251 | 18.74509804 | 0.018563723 |
| Acid_phosphat_B | 0.000561339 | 2.68320253 | 33.19444444 | 0.018633351 |
| Baculo_PP31 | 3.74E-05 | 3.74E-05 | 531.1111111 | 0.019852837 |
| PUD | 0.001143429 | 5.465590725 | 18.17490494 | 0.020781714 |
| LolA | 0.001143429 | 5.465590725 | 18.24427481 | 0.020861033 |
| EnY2 | 0.001143429 | 5.465590725 | 18.31417625 | 0.020940961 |
| CobD_Cbib | 0.001143429 | 5.465590725 | 18.38461538 | 0.021021503 |
| PAP2 | 0.001218857 | 5.82613772 | 17.96992481 | 0.021902773 |
| LON | 0.001210681 | 5.787057147 | 18.10606061 | 0.021920671 |
| LtrA | 0.001218857 | 0.001218857 | 18.03773585 | 0.021985425 |
| ASRT | 3.74E-05 | 0.178675532 | 597.5 | 0.022334441 |
| BPD_transp_1 | 0.001305068 | 6.238225717 | 17.90262172 | 0.023364141 |
| GARS_N | 0.001376842 | 6.581303965 | 17.8358209 | 0.024557104 |
| GSPII_F | 0.001404407 | 0.001404407 | 17.7037037 | 0.024863197 |
| Elong-fact-P_C | 0.001404407 | 0.001404407 | 17.76951673 | 0.024955625 |
| Ribosomal_60s | 0.001444611 | 0.001444611 | 17.44525547 | 0.025201605 |
| HVSL | 0.001444611 | 0.001444611 | 17.50915751 | 0.025293918 |
| Cad | 0.001444611 | 0.001444611 | 17.57352941 | 0.02538691 |
| Malectin_like | 0.001470114 | 0.001470114 | 17.31884058 | 0.025460668 |
| ATP-synt_8 | 0.001444611 | 0.001444611 | 17.63837638 | 0.025480589 |
| ArsD | 3.74E-05 | 0.178675532 | 682.8571429 | 0.025525076 |
| GATase_3 | 0.001470114 | 7.027144405 | 17.38181818 | 0.025553252 |
| VIT1 | 0.001547407 | 7.396606992 | 17.25631769 | 0.026702552 |
| Phg_2220_C | 0.001828411 | 8.739805985 | 14.79876161 | 0.027058223 |
| CBM_20 | 0.00183939 | 8.792283166 | 14.75308642 | 0.027136676 |
| Borrelia_REV | 0.001828411 | 0.001828411 | 14.8447205 | 0.027142255 |
| AFG1_ATPase | 0.001828411 | 8.739805985 | 14.89096573 | 0.02722681 |

| | | | | |
|---|---|---|---|---|
| YonK | 0.001828411 | 8.739805985 | 14.9375 | 0.027311894 |
| UxuA | 0.001828411 | 0.001828411 | 14.98432602 | 0.027397511 |
| Tom37 | 0.001828411 | 8.739805985 | 15.03144654 | 0.027483667 |
| TatC | 0.001828411 | 0.001828411 | 15.07886435 | 0.027570366 |
| Patched | 0.001648122 | 7.878023627 | 16.77192982 | 0.027642188 |
| SRCR | 0.001828411 | 8.739805985 | 15.12658228 | 0.027657614 |
| WW | 0.001647983 | 0.001647983 | 16.83098592 | 0.027737186 |
| SelA | 0.001828411 | 0.001828411 | 15.17460317 | 0.027745416 |
| RPA_C | 0.001668714 | 0.001668714 | 16.65505226 | 0.027792521 |
| Saf-Nte_pilin | 0.001828411 | 8.739805985 | 15.22292994 | 0.027833777 |
| Ribosomal_L37ae | 0.001647983 | 7.877360865 | 16.89045936 | 0.027835197 |
| Peptidase_C11 | 0.001906139 | 0.001906139 | 14.617737 | 0.027863438 |
| Peptidase_S8 | 0.001679072 | 0.001679072 | 16.59722222 | 0.027867926 |
| SBF | 0.001894909 | 0.001894909 | 14.70769231 | 0.027869744 |
| RNA_POL_M_15KD | 0.001668714 | 7.976453638 | 16.71328671 | 0.027889698 |
| RNA_pol_Rpb4 | 0.001828411 | 8.739805985 | 15.2715655 | 0.027922703 |
| MetJ | 0.001647983 | 7.877360865 | 16.95035461 | 0.027933904 |
| ApbE | 0.001906139 | 9.111344265 | 14.66257669 | 0.027948909 |
| Pox_ATPase-GT | 0.001828411 | 0.001828411 | 15.32051282 | 0.028012199 |
| Phage_GP20 | 0.00169991 | 0.00169991 | 16.48275862 | 0.028019207 |
| Endonuclease_NS | 0.001647983 | 7.877360865 | 17.01067616 | 0.028033313 |
| Phage_head_chap | 0.001828411 | 8.739805985 | 15.36977492 | 0.02810227 |
| Sex_peptide | 0.001928781 | 9.219572798 | 14.57317073 | 0.028108454 |
| AXE1 | 0.00169991 | 0.00169991 | 16.53979239 | 0.028116159 |
| Borrelia_orfA | 0.001647983 | 7.877360865 | 17.07142857 | 0.028133432 |
| Sigma54_DBD | 0.001637887 | 7.829102222 | 17.1942446 | 0.028162238 |
| Lectin_N | 0.001644054 | 7.858577209 | 17.13261649 | 0.028166943 |
| Pep_deformylase | 0.001828411 | 0.001828411 | 15.41935484 | 0.028192923 |
| Peptidase_M50 | 0.001731476 | 8.276457633 | 16.31399317 | 0.028247296 |
| Lipoprotein_11 | 0.001828411 | 0.001828411 | 15.46925566 | 0.028284162 |
| PAS_3 | 0.001731476 | 0.001731476 | 16.36986301 | 0.028344033 |
| Amidohydro_2 | 0.001951478 | 9.328064365 | 14.52887538 | 0.028352779 |
| Lact_bio_phlase | 0.001828411 | 8.739805985 | 15.51948052 | 0.028375993 |
| RskA | 0.001730382 | 8.271223912 | 16.42611684 | 0.02842345 |
| Hepar_II_III | 0.001962918 | 0.001962918 | 14.48484848 | 0.028432569 |
| IF3_C | 0.001828411 | 0.001828411 | 15.57003257 | 0.028468423 |
| H_kinase_N | 0.001828411 | 0.001828411 | 15.62091503 | 0.028561457 |
| HicB | 0.001828411 | 0.001828411 | 15.67213115 | 0.028655102 |
| HHH | 0.001828411 | 8.739805985 | 15.72368421 | 0.028749362 |
| GAF | 0.001828411 | 8.739805985 | 15.77557756 | 0.028844244 |
| NTP_transf_2 | 0.002009101 | 0.002009101 | 14.39759036 | 0.028926206 |
| Flg_hook | 0.001828411 | 8.739805985 | 15.82781457 | 0.028939755 |
| Semialdhyde_dh | 0.002020752 | 0.002020752 | 14.35435435 | 0.02900659 |
| GFA | 0.002009101 | 9.603500528 | 14.44108761 | 0.029013597 |
| eIF3g | 0.001828411 | 8.739805985 | 15.88039867 | 0.0290359 |
| GSHPx | 0.002044182 | 0.002044182 | 14.22619048 | 0.029080924 |

| | | | | |
|---|---|---|---|---|
| Arginase | 0.002032446 | 9.715091012 | 14.31137725 | 0.029087099 |
| EI24 | 0.001828411 | 8.739805985 | 15.93333333 | 0.029132687 |
| Cna_B | 0.002044182 | 9.771190419 | 14.26865672 | 0.029167733 |
| DHquinase_I | 0.001828411 | 0.001828411 | 15.98662207 | 0.02923012 |
| MACPF | 0.002086968 | 9.97570645 | 14.01759531 | 0.029254271 |
| Tn916-Xis | 0.002079646 | 0.002079646 | 14.10029499 | 0.029323621 |
| CsgG | 0.001828411 | 0.001828411 | 16.04026846 | 0.029328208 |
| Fer4_3 | 0.002067782 | 0.002067782 | 14.18397626 | 0.029329372 |
| ACCA | 0.002086968 | 9.97570645 | 14.05882353 | 0.029340313 |
| RseC_MucC | 0.002079646 | 9.94070745 | 14.14201183 | 0.029410377 |
| HEPN | 0.002154366 | 10.29787059 | 13.65714286 | 0.029422487 |
| CbiK | 0.001828411 | 8.739805985 | 16.09427609 | 0.029426956 |
| DNA_RNApol_7kD | 0.002151724 | 0.002151724 | 13.69627507 | 0.029470607 |
| GvpD | 0.002128669 | 0.002128669 | 13.85507246 | 0.029492862 |
| T4_deiodinase | 0.002142684 | 10.24202998 | 13.77521614 | 0.029515937 |
| Cas_Cas1 | 0.001828411 | 8.739805985 | 16.14864865 | 0.029526372 |
| AAA_2 | 0.002151724 | 10.28524191 | 13.73563218 | 0.029555293 |
| RPN7 | 0.002115493 | 0.002115493 | 13.97660819 | 0.029567416 |
| Glyco_hydro_38C | 0.002128669 | 10.17503727 | 13.89534884 | 0.029578597 |
| CheR | 0.002142684 | 10.24202998 | 13.8150289 | 0.029601243 |
| Methyltransf_2 | 0.001828029 | 0.001828029 | 16.20338983 | 0.029620267 |
| EVE | 0.002128669 | 0.002128669 | 13.93586006 | 0.029664832 |
| LGFP | 0.001828029 | 0.001828029 | 16.2585034 | 0.029721016 |
| Arrestin_N | 3.74E-05 | 3.74E-05 | 796.6666667 | 0.029779255 |
| PadR | 0.002206271 | 0.002206271 | 13.57954545 | 0.029960154 |
| HHH_2 | 0.002206271 | 10.54597407 | 13.61823362 | 0.03004551 |
| Wzz | 0.002225343 | 0.002225343 | 13.50282486 | 0.03004842 |
| SKN1 | 0.002225343 | 10.63714075 | 13.54107649 | 0.030133543 |
| Na_H_Exchanger | 0.002282637 | 0.002282637 | 13.46478873 | 0.030735221 |
| SLBB | 0.002351485 | 0.002351485 | 13.24099723 | 0.031136007 |
| Dna2 | 0.002348243 | 11.22460121 | 13.27777778 | 0.031179448 |
| OrfB_Zn_ribbon | 0.002364337 | 0.002364337 | 13.20441989 | 0.031219698 |
| CHB_HEX | 0.002345141 | 0.002345141 | 13.31476323 | 0.031225003 |
| Relaxase | 0.002338676 | 11.17887338 | 13.35195531 | 0.031225903 |
| Amino_oxidase | 0.002325911 | 11.11785521 | 13.42696629 | 0.03122993 |
| Glyco_hydro_92 | 0.002338676 | 0.002338676 | 13.38935574 | 0.031313371 |
| Phage_CI_repr | 0.002390171 | 11.42501664 | 13.16804408 | 0.031473875 |
| Thymidylate_kin | 0.002402044 | 0.002402044 | 13.13186813 | 0.031543325 |
| Glyco_hydro_65m | 0.002417317 | 11.55477291 | 13.09589041 | 0.031656912 |
| GFO_IDH_MocA | 0.00244236 | 0.00244236 | 13.06010929 | 0.031897485 |
| DnaB_C | 0.002455516 | 0.002455516 | 13.02452316 | 0.031981921 |
| ADH_zinc_N | 0.002497529 | 11.93818861 | 12.91891892 | 0.032265375 |
| CxxC_CxxC_SSSS | 0.002495245 | 11.92727115 | 12.95392954 | 0.032323228 |
| Baculo_PEP_C | 0.002495245 | 0.002495245 | 12.98913043 | 0.032411063 |
| CBM_5_12 | 0.002535367 | 12.11905428 | 12.88409704 | 0.032665914 |
| TraG-D_C | 0.002548828 | 0.002548828 | 12.84946237 | 0.032751075 |

| | | | | |
|---|---|---|---|---|
| UPF0122 | 0.002699796 | 12.90502518 | 12.16284987 | 0.032837214 |
| ChitinaseA_N | 0.002817002 | 13.46527062 | 11.65853659 | 0.032842123 |
| TPR_1 | 0.002812934 | 0.002812934 | 11.68704156 | 0.032874872 |
| Redoxin | 0.002699796 | 0.002699796 | 12.19387755 | 0.032920983 |
| PhageMin_Tail | 0.002713784 | 12.97188743 | 12.1319797 | 0.032923572 |
| HTH_18 | 0.002770176 | 13.24144228 | 11.89054726 | 0.032938911 |
| NAD_binding_4 | 0.002784385 | 13.309358 | 11.83168317 | 0.032943955 |
| HTH_AraC | 0.002798637 | 0.002798637 | 11.77339901 | 0.03294947 |
| STN | 0.002812934 | 0.002812934 | 11.71568627 | 0.032955447 |
| Poly_export | 0.002833653 | 13.54486046 | 11.63017032 | 0.032955865 |
| Glyco_tran_WecB | 0.002768315 | 13.23254471 | 11.9201995 | 0.032998865 |
| NHL | 0.002699796 | 12.90502518 | 12.22506394 | 0.033005179 |
| AhpC-TSA | 0.002575882 | 12.31271795 | 12.8150134 | 0.033009968 |
| Pyr_redox_2 | 0.002727816 | 0.002727816 | 12.10126582 | 0.033010026 |
| SNF2_N | 0.002756012 | 0.002756012 | 11.97994987 | 0.033016886 |
| ACP_syn_III | 0.002735297 | 0.002735297 | 12.07070707 | 0.033016968 |
| FAD_binding_3 | 0.002784385 | 13.309358 | 11.86104218 | 0.033025702 |
| DivIC | 0.002798637 | 0.002798637 | 11.80246914 | 0.033030826 |
| Hexapep | 0.002812934 | 13.44582255 | 11.74447174 | 0.033036419 |
| PrpR_N | 0.002767407 | 0.002767407 | 11.95 | 0.033070513 |
| TCL1_MTCP1 | 0.002691894 | 12.86725243 | 12.28791774 | 0.03307777 |
| N6_Mtase | 0.002699796 | 12.90502518 | 12.25641026 | 0.033089808 |
| Pyr_redox | 0.002756012 | 13.17373748 | 12.01005025 | 0.033099843 |
| SAND | 0.002691894 | 12.86725243 | 12.31958763 | 0.033163022 |
| HTH_19 | 0.002756012 | 0.002756012 | 12.04030227 | 0.033183218 |
| PMT | 0.002691894 | 0.002691894 | 12.35142119 | 0.033248714 |
| Melibiase | 0.002603112 | 0.002603112 | 12.78074866 | 0.033269715 |
| NMT_C | 0.002691894 | 0.002691894 | 12.38341969 | 0.033334851 |
| TonB | 0.002616792 | 12.50826551 | 12.74666667 | 0.033355375 |
| Cytochrom_CIII | 0.002691894 | 12.86725243 | 12.41558442 | 0.033421435 |
| Phage_integrase | 0.00288508 | 0.00288508 | 11.60194175 | 0.033472529 |
| Cytochrom_C552 | 0.002691894 | 0.002691894 | 12.44791667 | 0.03350847 |
| Sigma54_activat | 0.002671952 | 0.002671952 | 12.54593176 | 0.03352213 |
| Peptidase_S24 | 0.002644284 | 12.63967876 | 12.67904509 | 0.033526999 |
| RrnaAD | 0.002669818 | 0.002669818 | 12.57894737 | 0.033583496 |
| BCA_ABC_TP_C | 0.002691894 | 0.002691894 | 12.48041775 | 0.033595959 |
| T4SS-DNA_transf | 0.002685852 | 12.8383733 | 12.51308901 | 0.033608307 |
| N6_N4_Mtase | 0.002644284 | 0.002644284 | 12.71276596 | 0.033616167 |
| FlgM | 0.002669818 | 12.76172855 | 12.6121372 | 0.033672107 |
| Ni_hydr_CYTB | 0.002986473 | 0.002986473 | 11.30023641 | 0.033747849 |
| Ephrin | 0.002669818 | 0.002669818 | 12.64550265 | 0.033761187 |
| Mannitol_dh | 0.002986473 | 14.27533994 | 11.32701422 | 0.03382782 |
| MAAL_N | 0.002986473 | 0.002986473 | 11.35391924 | 0.033908171 |
| WRKY | 0.002985471 | 0.002985471 | 11.38095238 | 0.033977502 |
| UTRA | 0.002985471 | 0.002985471 | 11.40811456 | 0.034058594 |
| NAGLU | 0.002944259 | 14.07356008 | 11.57384988 | 0.034076417 |

| | | | | |
|---|---|---|---|---|
| Tup_N | 0.002985471 | 0.002985471 | 11.4354067 | 0.034140074 |
| HlyIII | 0.002985471 | 0.002985471 | 11.46282974 | 0.034221945 |
| Ketoacyl-synt_C | 0.002964966 | 14.17253513 | 11.54589372 | 0.034233177 |
| MFS_1_like | 0.003040633 | 0.003040633 | 11.27358491 | 0.034278833 |
| PNP_UDP_1 | 0.003056863 | 0.003056863 | 11.22065728 | 0.03430001 |
| ApoA-II | 0.002985471 | 0.002985471 | 11.49038462 | 0.034304209 |
| Glyco_hydro_98M | 0.003056863 | 14.61180416 | 11.24705882 | 0.034380716 |
| AdoMet_Synthase | 0.002985471 | 0.002985471 | 11.51807229 | 0.03438687 |
| TMV_coat | 0.003108171 | 14.85705895 | 11.1682243 | 0.034712755 |
| HTH_1 | 0.003108171 | 14.85705895 | 11.19437939 | 0.034794049 |
| TrbC | 0.003150711 | 0.003150711 | 11.14219114 | 0.035105822 |
| OmpA | 0.003198606 | 0.003198606 | 11.11627907 | 0.035556593 |
| Apo-CII | 3.74E-05 | 0.178675532 | 956 | 0.035735106 |
| HxlR | 0.003317282 | 15.85660905 | 11.09048724 | 0.036790276 |
| Sigma70_r4 | 0.003394507 | 16.22574546 | 11.06481481 | 0.037559596 |
| Rab5-bind | 0.003494133 | 0.003494133 | 11.03926097 | 0.038572643 |
| TBPIP | 0.003528749 | 0.003528749 | 11.01382488 | 0.038865028 |
| FAD_binding_2 | 0.003725869 | 0.003725869 | 10.98850575 | 0.04094173 |
| DUF3876 | 0.003744101 | 0.003744101 | 10.96330275 | 0.041047709 |
| IFT46_B_C | 0.003801888 | 0.003801888 | 10.9382151 | 0.041585867 |
| Esterase_phd | 0.003861759 | 18.45920877 | 10.86363636 | 0.041952747 |
| Thioredoxin | 0.003854585 | 0.003854585 | 10.88838269 | 0.041970201 |
| CtnDOT_TraJ | 0.003873182 | 0.003873182 | 10.83900227 | 0.041981426 |
| COG2 | 0.003854585 | 18.42491809 | 10.91324201 | 0.042066023 |
| GDPD | 0.003922176 | 0.003922176 | 10.81447964 | 0.042416289 |
| NAGLU_C | 0.003971383 | 0.003971383 | 10.79006772 | 0.042851493 |
| Fer4_7 | 0.004004815 | 0.004004815 | 10.76576577 | 0.0431149 |
| HTH_11 | 0.004042894 | 19.32503123 | 10.74157303 | 0.043427036 |
| TRCF | 0.004197293 | 20.06306118 | 10.57522124 | 0.044387303 |
| OSTMP1 | 0.004197293 | 0.004197293 | 10.59866962 | 0.044485723 |
| DPM3 | 0.004222025 | 0.004222025 | 10.55187638 | 0.04455029 |
| NlpE | 0.004169296 | 19.92923646 | 10.6935123 | 0.044584422 |
| LIP | 0.004197293 | 0.004197293 | 10.62222222 | 0.04458458 |
| Antimicrobial12 | 3.74E-05 | 3.74E-05 | 1195 | 0.044668883 |
| HasA | 0.004197293 | 0.004197293 | 10.64587973 | 0.044683878 |
| Glu_cyclase_2 | 0.004169296 | 19.92923646 | 10.71748879 | 0.044684387 |
| Mannitol_dh_C | 0.004197293 | 0.004197293 | 10.66964286 | 0.044783619 |
| SRR | 0.004282057 | 0.004282057 | 10.52863436 | 0.045084208 |
| N_NLPC_P60 | 0.004360367 | 20.84255451 | 10.43668122 | 0.045507761 |
| Lambda_Bor | 0.004360367 | 0.004360367 | 10.4595186 | 0.04560734 |
| Bro-N | 0.0043819 | 20.94548374 | 10.41394336 | 0.045632862 |
| ABC_membrane | 0.004360367 | 0.004360367 | 10.48245614 | 0.045707356 |
| MarR | 0.004354207 | 20.81311061 | 10.50549451 | 0.0457431 |
| Resolvase | 0.004500361 | 0.004500361 | 10.39130435 | 0.046764624 |
| Phage_Mu_F | 0.00455635 | 0.00455635 | 10.32397408 | 0.047039637 |
| NTP_transferase | 0.00455635 | 21.77935206 | 10.34632035 | 0.047141455 |

| | | | | |
|---|---|---|---|---|
| PDDEXK_2 | 0.004555621 | 0.004555621 | 10.36876356 | 0.047236155 |
| Response_reg | 0.004606443 | 22.01879529 | 10.30172414 | 0.0474543 |
| Ribosomal_L32p | 0.004741075 | 22.66233679 | 10.08438819 | 0.047810837 |
| Steroid_dh | 0.004734865 | 0.004734865 | 10.10570825 | 0.047849163 |
| Peptidase_S51 | 0.004734865 | 22.63265403 | 10.12711864 | 0.047950538 |
| FBPase_2 | 0.004734865 | 22.63265403 | 10.14861996 | 0.048052344 |
| Glyco_hydro_3_C | 0.004779867 | 22.84776264 | 10.06315789 | 0.048100553 |
| HTH_3 | 0.004713928 | 0.004713928 | 10.21367521 | 0.048146527 |
| DNA_topoisoIV | 0.004734865 | 22.63265403 | 10.17021277 | 0.048154583 |
| Glyoxalase | 0.004689733 | 0.004689733 | 10.27956989 | 0.048208437 |
| Fer4_2 | 0.004713928 | 22.53257449 | 10.23554604 | 0.048249624 |
| Abhydrolase_4 | 0.004734865 | 22.63265403 | 10.19189765 | 0.048257258 |
| Bac_transf | 0.004706963 | 0.004706963 | 10.25751073 | 0.048281727 |
| Glyco_hydro_10 | 0.004825297 | 0.004825297 | 10.04201681 | 0.048455713 |
| Lipocalin_3 | 0.005149067 | 0.005149067 | 9.42800789 | 0.048545445 |
| RRM_3 | 0.005144851 | 24.59238718 | 9.446640316 | 0.048601556 |
| Regulator_TrmB | 0.005144851 | 0.005144851 | 9.465346535 | 0.048697796 |
| MFS_1 | 0.005221222 | 24.95744288 | 9.3359375 | 0.048745006 |
| YadA | 0.005307905 | 0.005307905 | 9.192307692 | 0.0487919 |
| PspC | 0.005144851 | 0.005144851 | 9.484126984 | 0.048794419 |
| RNA12 | 0.005219108 | 0.005219108 | 9.354207436 | 0.048820617 |
| MatE | 0.005245056 | 0.005245056 | 9.317738791 | 0.04887206 |
| PYNP_C | 0.005307905 | 0.005307905 | 9.210019268 | 0.048885912 |
| Proteasom_PSMB | 0.005144851 | 24.59238718 | 9.502982107 | 0.048891426 |
| Ogr_Delta | 0.005219108 | 0.005219108 | 9.37254902 | 0.048916343 |
| OAD_gamma | 0.005307905 | 25.37178809 | 9.227799228 | 0.048980286 |
| Prenyltransf | 0.005144851 | 0.005144851 | 9.521912351 | 0.048988819 |
| Beta_elim_lyase | 0.005206951 | 0.005206951 | 9.409448819 | 0.048994542 |
| HGD-D | 0.005219108 | 24.94733507 | 9.390962672 | 0.049012446 |
| CUT | 0.005307905 | 25.37178809 | 9.245647969 | 0.049075025 |
| PGP_phosphatase | 0.005144851 | 24.59238718 | 9.540918164 | 0.049086601 |
| Cenp-O | 0.005307905 | 25.37178809 | 9.263565891 | 0.049170132 |
| PC4 | 0.005144851 | 0.005144851 | 9.56 | 0.049184774 |
| SusD-like_2 | 0.005292904 | 25.30007901 | 9.299610895 | 0.049221944 |
| Brevenin | 0.005307905 | 0.005307905 | 9.281553398 | 0.049265608 |
| Pam17 | 0.005144851 | 0.005144851 | 9.579158317 | 0.049283341 |
| NRPS | 0.005144851 | 24.59238718 | 9.598393574 | 0.049382304 |
| Myf5 | 0.005144851 | 0.005144851 | 9.617706237 | 0.049481664 |
| Glyco_hydro_18 | 0.005397876 | 25.80184915 | 9.174664107 | 0.049523703 |
| MRP-S27 | 0.005144851 | 24.59238718 | 9.637096774 | 0.049581426 |
| MOFRL | 0.005144851 | 0.005144851 | 9.656565657 | 0.04968159 |
| MmgE_PrpD | 0.005144851 | 0.005144851 | 9.67611336 | 0.04978216 |
| Lipoprotein_X | 0.005144851 | 0.005144851 | 9.695740365 | 0.049883138 |
| Myosin_tail_1 | 0.00544842 | 0.00544842 | 9.157088123 | 0.049891661 |
| LIM_bind | 0.005144851 | 24.59238718 | 9.715447154 | 0.049984527 |
| HRDC | 0.005144851 | 24.59238718 | 9.735234216 | 0.050086328 |

| | | | | |
|---|---|---|---|---|
| Herpes_UL6 | 0.005144851 | 0.005144851 | 9.755102041 | 0.050188545 |
| Hemopexin | 0.005144851 | 0.005144851 | 9.775051125 | 0.05029118 |
| TarH | 0.005038417 | 24.08363323 | 10 | 0.05038417 |
| Glyco_hydro_63 | 0.005144851 | 0.005144851 | 9.795081967 | 0.050394236 |
| Intron_maturas2 | 0.005032728 | 24.05644196 | 10.02096436 | 0.050432792 |
| FliS | 0.005144851 | 0.005144851 | 9.815195072 | 0.050497715 |
| FlgI | 0.005144851 | 0.005144851 | 9.835390947 | 0.05060162 |
| DSPc | 0.005144851 | 0.005144851 | 9.855670103 | 0.050705953 |
| DNA_ligase_A_N | 0.005144851 | 24.59238718 | 9.876033058 | 0.050810717 |
| DIL | 0.005144851 | 0.005144851 | 9.896480331 | 0.050915915 |
| CcmD | 0.005144851 | 0.005144851 | 9.917012448 | 0.05102155 |
| Calsequestrin | 0.005144851 | 24.59238718 | 9.937629938 | 0.051127624 |
| Antimicrobial20 | 0.005144851 | 0.005144851 | 9.958333333 | 0.05123414 |
| 3A | 0.005144851 | 24.59238718 | 9.979123173 | 0.051341101 |
| Anemone_cytotox | 3.74E-05 | 3.74E-05 | 1593.333333 | 0.059558511 |
| AlaDh_PNT_N | 3.74E-05 | 0.178675532 | 2390 | 0.089337766 |
| AA_kinase | 3.74E-05 | 3.74E-05 | 4780 | 0.178675532 |
