## Supplementary table 3 for "Punchline: Identifying and comparing significant Pfam protein domain differences across draft whole genome sequences"

| Pfam domain name | Suggested function conferred [1,2,3,4] |
|---|---|
| AAT | Cysteine peptidase |
| AA_kinase | Amino acid kinase |
| AlaDh_PNT_N | Alanine dehydrogenase/pyridine nucleotide transhydrogenase |
| Anemone_cytotox | Basic cytolysin |
| Antimicrobial12 | Antimicrobial peptide |
| Apo-CII | Activator for lipoprotein lipase |
| Arrestin_N | Ig-like beta-sandwich, e.g. thioredoxin-interacting protein |
| ArsD | Detoxification of arsenicals or general trans-acting repressor protein |
| ASRT | Small molecule binding domain |
| B12-binding_2 | Domain binds cobalamin (vitamin B12) |
| Baculo_PP31 | Possible phage domain |
| Bac_luciferase | Flavin monooxygenase |
| Bac_surface_Ag | Bacterial surface antigen |
| Borrelia_P13 | Integral membrane protein domain with surface-exposed regions |
| BPL_N | Unknown structural domain |
| Caa3_CtaG | Cytochrome oxidase caa3 type assembly factor |
| CAP_C | Adenylate cyclase associated C-terminal |
| CbiC | Cobalamin synthesis C-terminal |
| CbiG_mid | Anaerobic cobalamin synthesis mid domain |
| CbiG_N | Anaerobic cobalamin synthesis N-terminal |
| CHASE4 | Extracellular sensory domain |
| Coat_F | Bacillus spore-coat like domain |
| CobT | Anaerobic cobalt chelatase subunit for Vitamin B12 cobalamin synthesis |
| COG7 | Glycoconjugate synthesis |
| CRISPR_Cas2 | CRISPR-Cas domain |
| Cuticle_1 | Repeat protein domain |
| Desulfoferrod_N | Desulfoferrodoxin N-terminal domain |
| DmpG_comm | Aldolase C-terminal alpha helical domain |
| DsrC | Assembly, folding or stablilisation of siroheam proteins such as sulphite reductase and nitrite reductase |
| DUF605 | Domain of unknown function |
| eIF2A | Beta-propellor domain |
| ESR1_C | Receptor-like protein domain |
| Euplotes_phero | Protoxoan signalling pheromone domain |
| FAD_syn | Riboflavin kinase/FAD synthetase |
| FctA | Surface protein antigen |
| Flagellin_D3 | Flagellin central domain |
| Flavi_NS1 | Cysteine-rich putative phage protein |
| Flavokinase | Riboflavin/FAD synthetase C-terminal |
| Fucokinase | Fucose kinase |
| Fun_ATP-synt_8 | Transmembrane ATPase stalk-like membrane anchor |
| Gag_p24 | Putative phage domain |
| GCR1_C | Transcriptional activator |
| Glycogen_syn | Glycogen synthesis domain |
| Glyco_hydro_15 | Glycoside hydrolase family 15 |
| Glyco_hydro_38 | Glycoside hydrolase family 38 |
| Glyco_hydro_71 | Glycoside hydrolase family 71 |
| Gypsy | Putative phage domain |
| Herpes_LMP2 | Membrane protein domain, possible phage origin |
| HyaE | Hydrogenase-assembly like domain |
| IBR | Cysteine rich domain |
| Ion_trans_N | Na/K ion channel N-terminal |
| IP_trans | Phospholipid transport |
| LAM_C | Lysine-2,3-aminomutase |
| Lipase | Lipase |
| Lipase_bact_N | Lipase N terminal |
| Lon_C | Lon protease C-terminal |
| Malectin_like | Membrane-anchored domain binding glycans |
| MAP7 | Microtubule-stablising-like domain |
| Met_synt_B12 | Vitamin B12 dependent methionine synthase |
| MHCassoc_trimer | Antigen-like domain |
| Mig-14 | Antimicrobial peptide resistance |
| MrpF_PhaF | Na/K efflux regulation of pH |
| Mto1_bdg | Microtubule-stablising-like domain C-terminal |
| Nitro_FeMo-Co | Iron-Molybdenum cofactor/ribonuclease H-like domain |
| OmpW | Outer membrane W |
| Opacity | Porins related to OmpA-like domains |
| OpuAC | Osmoregulation high affinity transporter |
| P-II | Signalling protein in nitrogen metabolism |
| PEP-utilisers_N | Swivelling domain beta/beta/alpha |
| Peptidase_C37 | Serine/Cysteine protease |
| Phytoreo_S7 | Peptidase |
| Pkip-1 | Protein kinase-interacting domain |
| Pox_E2-like | Domain like pox E2 viral |
| PRELI | Globular alpha + beta fold |
| Prenyltrans | Repeat domain such as in prenyltransferase |
| PriCT_1 | Alpha-helical domain, primase-like |
| PS_Dcarbxylase | Phosphatidylserine decarboxylase |
| PulG | Pseudopilin |
| Pyridox_ox_2 | Pyridoxamine 5'phosohate oxidase-related |
| QRPTase_N | Quinolinate phosphoribosyl transferase/nicotinate nucleotide pyrophosphorylase |
| Rap1_C | All helical fold telomeric-like domain |
| RNA_ligase | RNA ligases in DNA-RNA repair |
| S10_plectin | Potential RNA-binding domain |
| SAPS | Associating with phosphatase |
| SBP_bac_3 | Bacterial extracellular solute domain |
| Selenoprotein_S | Peroxidase and reductase selenoprotein-like |
| SLH | Non-covalently anchored to cell surface via wall polysaccharide pyruvylation |
| SpoIIP | Autolysin with peptidoglycan hydrolase activity |
| SURNod19 | Stress upregulated Nod-like |
| TauE | Sulfite exporter |
| Thg1 | RRM ferredoxin fold palm domain, like polymerase of CRISPR, polymerases and diguanylate cyclases |
| TIR-like | Nucleotide binding domain |
| TPK_catalytic | Thiamine pyrophosphokinase |
| TRAP-gamma | Translocon-associated protein |
| Uteroglobin | Small disulphide-bridged mammalian goblin-like |
| Wax2_C | Short chain dehydrogenase C-terminal |
| YLP | Unknown function, possible tyrosine kinase substrate |
| YMF19 | Transmembrane ATPase stalk-like membrane anchor |
| zf-DNA_Pol | DNA/RNA/protein/lipid binding, Zn binding |
| zf-piccolo | DNA/RNA/protein/lipid binding, Zn binding |
| Zn_Tnp_IS1 | Insertion sequence element, Zn binding |


1. Finn, R. D. *et al.* InterPro in 2017-beyond protein family and domain annotations. *Nucleic Acids Res.* **45,** D190–D199 (2017).
2. Mitchell, A. L. *et al.* InterPro in 2019: Improving coverage, classification and access to protein sequence annotations. *Nucleic Acids Res.* **47,** D351–D360 (2019).
3. Eddy, S. R. Accelerated profile HMM searches. *PLoS Comput. Biol.* **7,** (2011).
4. El-Gebali, S. *et al.* The Pfam protein families database in 2019. *Nucleic Acids Res.* (2018). doi:10.1093/nar/gky995
