## Supplementary table 1 for "Punchline: Identifying and comparing significant Pfam protein domain differences across draft whole genome sequences"

Supplementary table 1
Accession numbers for genomes used in the study

### ***Bacteroides ovatus***

| Strain | Site | Notes | Isolate Name | Citation |
|---|---|---|---|---|
| GCF_000154125.1_ASM15412v1 | Lab(human) | lab | ovatus_ATCC8483 | Fulton et al., 2007, Wash. U |
| GCF_000178275.1_ASM17827v1 | human | healthy | ovatus_SD_CMC_3f | Nelson et al., 2010, JCVI |
| GCF_000218325.1_Bact_ovat_3_8_47FAA_V1 | human | Healthy | ovatus_3_8_47FAA | HMP reference genome, 2011, Broad Inst. |
| GCF_000273195.1_Bact_ovat_CL02T12C04_V1 | human | healthy | ovatus_CL02T12C04 | HMP reference genome, 2012, Broad Inst. |
| GCF_000273215.1_Bact_ovat_CL03T12C18_V1 | human | healthy | ovatus_CL03T12C18 | HMP reference genome, 2012, Broad Inst. |
| GCF_000699665.1_ASM69966v1 | human | ?disease | ovatus_3725-D9-iii | 2014, medical, U Maryland |
| GCF_000699725.1_ASM69972v1 | human | ?disease | ovatus_3725_D1-iv | 2014, medical, U Maryland |
| GCF_001314995.1_ASM131499v1 | human | healthy | ovatus_ATCC8483 | Wu et al., 2015 Science ,Wash U |
| GCF_001405735.1_14207_7_66 | human | healthy | ovatus_2789STDY5834943 | Browne et al., 2016, Nature |
| GCF_001535615.1_ASM153561v1 | human | ?disease | ovatus_CL09T03C03 | Coyne et al.,2016,Brigham& women's hospital |
| GCF_001578575.1_ASM157857v1 | Human | healthy | ovatus_KLE1656 | HMP reference strain, 2016, Wash U |
| GCF_900095495.1_B_ovatus_V975 | Human | healthy | ovatus_V975 | IFR, 2016 |
| GCF_900100465.1_IMG-taxon_2654588146 | cow | healthy | ovatus_NLAE-zl-C57 | JGI, 2016, Hungate 1000, cow faeces |
| GCF_900102645.1_IMG-taxon_2654588143 | cow | healthy | ovatus_NLAE-zl-C500 | JGI, 2016, Hungate 1000, cow faeces |
| GCF_900107475.1_IMG-taxon_2693429857 | lab | NA | ovatus_DSM1896 | JGI, 2016 |

### ***Bacteroides thetaiotaomicron***

| Strain | Site | Notes | Isolate Name | Citation |
|---|---|---|---|---|
| GCF_000011065.1_ASM1106v1 | human | healthy | thetaiotaomicron_VPI-5482 | Xu et al., 2003, Science, Bethesda |
| GCF_000304195.1_Bact_thet_CL09T03C10_V1 | human | healthy | thetaiotaomicron_CL09T03C10_V1_finegoldii | Ribeiro et al., 2012. Genome Res, Broad |
| GCF_000403155.2_Bact_thet_dnLKV9_V1 | human | healthy | thetaiotaomicron_dnLKV9 | Earl et al., 2013, Broad |
| GCF_001049535.1_3731 | human | healthy | thetaiotaomicron_3731 | Planer J, 2015, Washington U |
| GCF_001055755.1_ASM105575v1 | human | ?disease | thetaiotaomicron_19_BTHE | Roach et al., 2015, PLOS Gen. |
| GCF_001314975.1_ASM131497v1 | human | healthy | thetaiotaomicron_7330 | Wu et al., 2015, Science, Wash U |
| GCF_001373135.1_2e6A_assembly | human | undernourished | thetaiotaomicron_2e6A | Kau, A, 2015, Washington U |
| GCF_001405095.1_13470_2_65 | human | healthy | thetaiotaomicron_2789STDY5834846 | Pathogen Informatics, 2015, WTSI |
| GCF_001405255.1_13414_6_57 | human | healthy | thetaiotaomicron_2789STDY5608873 | Pathogen informatics, 2015, WTSI |
| GCF_001578565.1_ASM157856v1 | human | healthy | thetaiotaomicron_KLE1254 | Mitreva et al., 2016, Wash U |
| GCF_001816245.1_ASM181624v1 | human | ?disease | thetaiotaomicron_14-106904-2 | Sydenham et al.,2016, Odense, DK |
| GCF_900109385.1_IMG-taxon_2593339226 | Cow | healthy | thetaiotaomicron_KPPR-3 | Verghese & Submission, JGI, 2016 |

?disease - Individual host was in hospital or exhibited a disease
